# Vascular effects on the BOLD response and the retinotopic mapping of hV4

**DOI:** 10.1101/413492

**Authors:** H.G. Boyd Taylor, A.M. Puckett, Z.J. Isherwood, M.M. Schira

## Abstract

Despite general acceptance that the retinotopic organisation of human V4 (hV4) takes the form of a single, uninterrupted ventral hemifield, measured retinotopic maps of this visual area are often incomplete. Here, we test hypotheses that artefact from draining veins close to hV4 cause inverted BOLD responses that may serve to obscure a portion of the lower visual quarterfield — including the lower vertical meridian — in some hemispheres. We further test whether correcting such responses can restore the ‘missing’ retinotopic coverage in hV4. Subjects (N=11) viewed bowtie, ring, drifting bar and full field flash stimuli. Functional EPIs were acquired over approximately 1.5h and analysed to reveal retinotopic maps of early visual cortex, including hV4. Normalised mean maps (which show the average EPI signal amplitude) were constructed by voxel-wise averaging of the EPI time course and used to locate venous eclipses, which can be identified by a decrease in the EPI signal caused by deoxygenated blood. Inverted responses are shown to cluster in these regions, and correcting these responses improves maps of hV4 in some hemispheres, including restoring a complete hemifield map in one. A leftwards bias was found in which 11/11 hV4 maps in the left hemisphere were classified as incomplete, while this was the case in only 3/11 right hemisphere maps. Incomplete hV4 maps did not correspond with venous artefact in many instances, with incomplete maps being present in the absence of a venous eclipse and complete maps coexisting with a proximate venous eclipse. We also show that mean maps of upper surfaces (near the boundary between cortical grey matter and CSF) provide highly detailed maps of veins on the cortical surface. Results suggest that venous eclipses and inverted voxels can explain some incomplete hV4 maps, but cannot explain the remainder nor the leftwards bias in hV4 coverage reported here.

## Introduction

Human visual cortex is comprised of multiple orderly visual areas which are functionally and anatomically distinct from each other [1–4]. Many of these areas are organised according to the principle of retinotopy, which is to say they are topographically organised such that the spatial organisation of the retina, and therefore the visual field, is mapped to the neurons of each area [5–7]. These retinotopic maps are highly preserved across individuals and species, and hence the cortical organisation of visual areas is unlikely to be random, but an integral part of visuo-cortical function [8–10]. It is therefore important to determine the organisation and position of visual areas, as these factors are central to our understanding of visual information processing.

Functional magnetic resonance imaging (fMRI) is currently one of the most well-suited and popular tools for studying visual areas in human visual cortex, due to its safe and non-invasive nature; however, it is not without issue. The impact of veins on the measured blood-oxygen-level dependent (BOLD) response in fMRI has been emphasised as one such issue, both inside and outside the visual cortex [11–15]. Here, the strong paramagnetic effect of deoxyhaemoglobin (dHb) in veins changes the homogeneity of the adjacent local magnetic field, causing a reduction or loss of the fMRI signal in its vicinity [16, 17]. This hinders the usefulness of fMRI for studying some visual areas, as the BOLD signal can be obfuscated by venous artefact. For example, this artefact has been accused of contributing to the deterioration of fMRI measurements in human ventral occipital cortex, particularly in area V4 — the organisation of which remains unsettled despite years of experimental investigation and debate [15, 19–25].

It is arguable that the observance of ‘incomplete’ V4 maps on the ventral surface indicates that human V4 follows the same non-symmetric dorsal/ventral split seen in macaque V4; however, the evidence for a dorsal component of V4 in humans is limited [22, 24, 26]. Rather, these incomplete maps are likely a result of the proximity of V4 to the Transverse Sinuses (TSs), a pair of large veins that drain dHb from the back of the head [15, 18]. In 2010, Winawer and colleagues [15] demonsrated how surface veins, in particular the TS, can be a source of measurement insufficiency affecting hV4 maps. Veins are known to cause a lower mean intensity of voxels in their proximity due to the high concentrations of dHb present within them [11–15]. This phenomenon has been called the ‘venous eclipse’, and it offers an elegant explanation for incomplete hV4 maps, as the TS is anatomically close to this visual area [15]. Hence, it is likely that human V4 is organised in a single, continuous hemifield adjacent to ventral V3 [25] (hereafter called ‘hV4’, in accordance with accepted nomenclature for this model) and that incomplete hV4 maps are due to insufficiencies in measurement rather than being genuine portrayals of the existing retinotopic map [15, 27, 28]. A comparison of these two models is shown in Fig 1.

**Fig 1.**
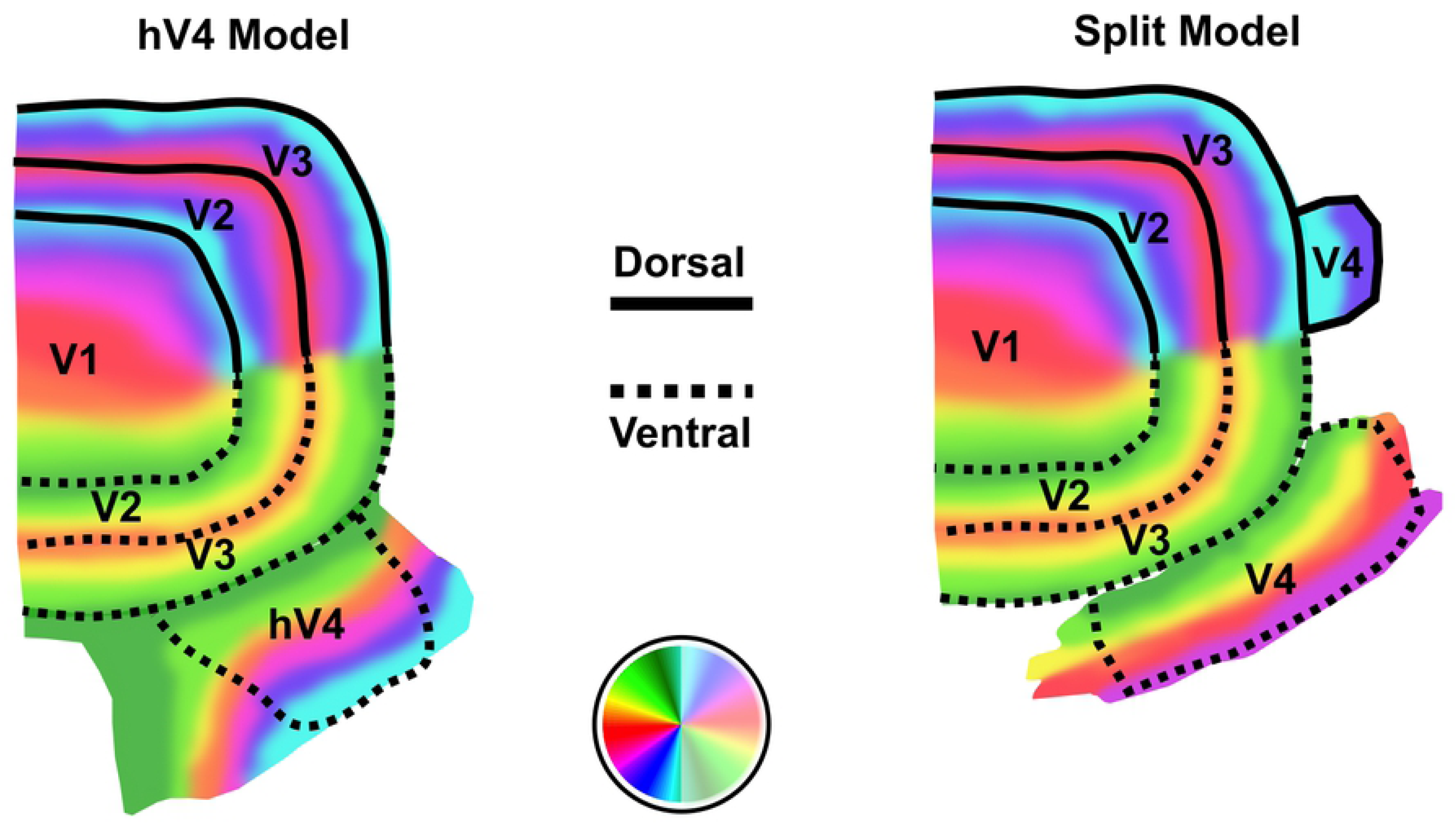
Models of human V4. Diagrams of early visual cortex in humans, showing the ‘hV4’ model (left), and the ‘Split’ model (right). According to the hV4 model, V4 contains a map of the entire contralateral visual hemifield in a continuous retinotopic map on the ventral surface, adjacent to ventral V3 (V3v). In this model, the lower boundary of hV4 is defined by a polar angle reversal at the lower vertical meridian on the ventral surface. The hV4 map illustrated here is based on descriptions provided in Winawer and Witthoft [28]. The ‘split’ model assumes the upper vertical meridian, contralateral horizontal meridian and approximately half the upper visual quarterfield are represented on the ventral surface, and the remainder of the map, including the lower vertical meridian is represented on the dorsal surface. The V4 map illustrated here is based on V4d and V4v retinotopic maps defined in Hansen and colleagues [22].

According to Winawer and colleagues [15], if artefact is near enough to the lower hV4 boundary, the measured map of hV4 will be incomplete due to signal dropout and, importantly, delays in the BOLD response. These delays resulted in the BOLD response being out-of-phase with the stimulus, being significantly delayed or even appearing counterphase to the stimulus, where voxels peak 180° later than in-phase voxels thus appearing between stimulus cycles rather than at the usual BOLD delay of approximately 6s [29]. However, these authors also note that when the TS was displaced by more than 1cm from the lower boundary of hV4, responses up to, or very close to the lower vertical meridian (i.e. complete maps) can be recorded [15].

Counterphase delays in the measured BOLD response along the lower hV4 boundary are one interpretation of the disrupted time courses of voxels affected by venous artefact, however another alternative exists. As the BOLD signal is driven by changes in oxygen levels in blood [17, 30], with oxygenated blood producing positive BOLD responses (PBRs), veins draining deoxygenated blood would theoretically result in a decrease in the mean intensity of nearby voxels. Therefore, the haemodynamic response of ‘venous’ voxels was proposed by Puckett and colleagues [31] to be more accurately described as in-phase but ‘inverted’ rather than counterphase.

In their study, Puckett and colleagues [31] correlated voxel responses to a full field control stimulus with a reference haemodynamic waveform. Ordinarily, the BOLD response is coupled with neural firing, therefore, the simultaneous stimulation of the whole visual field should result in all the voxels within the range of the stimulus exhibiting PBRs [15, 31], however many do not. Using this stimulus, Puckett and colleagues demonstrated many voxels in the region of the venous eclipse exhibit an inverted response to full field stimulation, rather than a delayed response [31]. They further showed that in early visual areas V1, V2 and V3, time courses of voxels exhibiting this response have been successfully corrected to restore the order of visual field maps. Whilst the same procedure was not applied to inverted voxels in hV4, this area was shown to contain a significantly higher number of inverted voxels compared with other areas [31]. This suggests that correcting inverted voxels in hV4 may enable a more accurate measure of its response in the region of the venous eclipse than has previously been possible.

Here, we aim to investigate the consistency of the venous eclipse and inverted voxel hypotheses proposed by Winawer and colleagues [15] and Puckett and colleagues [31], with relation to retinotopic maps of hV4. Based on the V4 literature in humans, we hypothesised that we would identify complete hV4 maps on the ventral surface of some hemispheres and incomplete maps in others. Furthermore, when an incomplete map was identified we hypothesised that a cause should be found, namely that a venous eclipse would also be present on or near the lower boundary of hV4, consistent with the hypothesis of Winawer and colleagues [15]. Additionally, it was predicted that in the region of the venous eclipse, voxels would present with on-time but inverted BOLD responses, as proposed by Puckett and colleagues [31]. Correcting these responses was predicted to result in the subsequent restoration of hV4 maps where they were affected by inverted voxels, potentially allowing a complete hemifield map to be identified where an incomplete map was initially measured. We further propose that any anomalous voxel responses, that is inverted, delayed or random responses, should be identifiable using a full field flash stimulus, as this should serve as a control that will distinguish voxels that fail to exhibit the expected BOLD response.

## Materials and methods

### Subjects

11 healthy subjects (5 female), aged 21-26 years with normal or corrected-to-normal vision participated in the study. The experiment was conducted with the understanding and written consent of each subject and was approved by the University of New South Wales Human Research Advisory Panel (HC14262).

### Visual field mapping

Travelling wave stimuli in the form of rotating bowties and expanding rings were used, adhering to the methods described in previous work [32, 33]. Stimuli were composed of a flickering chequerboard pattern under an extended grey fixation grid, with colours randomly changing every 0.25ms. A 3×3 pixel fixation dot (0.04°) was present in the centre of the screen, which changed colour every 3-8s, with subjects being instructed to press and hold a button whenever this dot was red. The bowties were presented for 15 cycles per scan, and the rings for 12.

Two additional types of stimuli were used to enable additional analyses. The first of these was a full field flash which was composed of the same flickering chequerboard pattern and fixation grid as the bowties and rings. It was presented for 12 cycles per scan, where each cycle was comprised of an ON period of 4s and an OFF period of 16s. The other stimulus was a drifting bar, adapted from the same stimulus used by Dumoulin and Wandell [34], with the addition of a grey fixation grid [33]. The bar was composed of two black and white chequered bars, one inside the other, coasting in opposite directions. The bar was presented at four orientations (0°, 45°, 90° and 115°), and travelled in the two directions perpendicular to each, for a total of eight directions. Each bar sweep took 40s and there were four periods of a blank luminance block, each lasting 20s, inserted between every second sweep (Fig 2).

**Fig 2.**
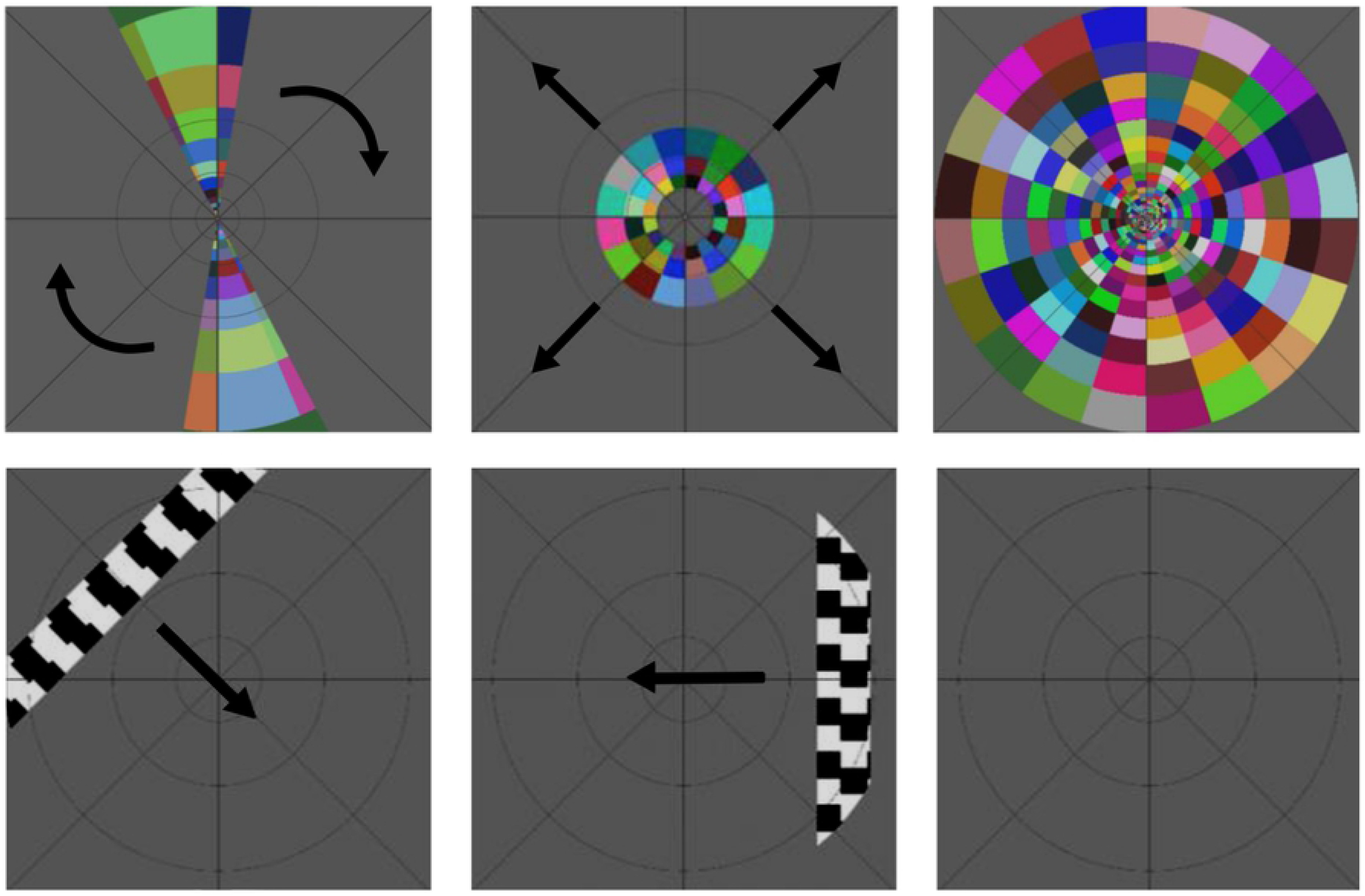
Stimuli. Top Row: Clockwise rotating bowtie, expanding rings and full field flash. Bottom Row: Drifting bar stimulus at three different time points, showing two of the eight bar directions and a blank luminance block. Arrows indicate the direction of movement.

### Stimulus presentation

Stimuli were generated using MATLAB R2012a and Psychophysics Toolbox [35, 36], and displayed to participants on a 19” monitor with a 1024 × 768mm resolution. The participants viewed this monitor via a mirror attached to the head coil. The stimuli subtended a visual display spanning 5.5° eccentricity (or 11° of visual angle), with a viewing distance of 1.5m. Total stimuli presentation and scan time per subject varied, with 3 subjects viewing only one presentation of each stimulus, resulting in a viewing time of approximately 24m, and the remaining 8 subjects viewing each stimulus up to 4 times, resulting in total viewing time of 1.5-2h. For 5 of these subjects, data collection was split across two scanning sessions, one of 25m and the other 1h; the 3 remaining subjects were scanned in one 2h session.

### Data acquisition

Functional EPI data was collected using a Philips 3T Achieva TX Magnetic Resonance Scanner fitted with Quasar Dual gradients and a 32-channel head coil. 32 oblique, ascending coronal slices covering the occipital pole and ventral occipital cortex were acquired at a voxel resolution of 1.5*mm*^3^, a 128 × 128 matrix and a field of view (FOV) of 192mm. Images were acquired with a repetition time of 2s, echo time of 25ms and acceleration (SENSE) factor of 2. Higher resolution T1 anatomical images were collected at a resolution of 0.75*mm*^3^, using a 3D magnetisation-prepared rapid acquisition with gradient echo (MPRAGE) protocol with a matrix size of a 340 × 340, a FOV of 256mm and a TR of 6s. The number of volumes collected varied by stimulus. Bowties, rings and full field scans had some initial volumes discarded to compensate for the initial pulse of the scanner and enable the BOLD response to reach baseline. Details of volumes are as follows — Bowties: 186 volumes, discarded first 6; Rings: 174 volumes, discarded first 6; Full field: 124 volumes, discarded first 4. Zero volumes were discarded from the bars as this stimulus began with an extended baseline (blank luminance block and fixation grid) of 20s. A venogram from Subject 4 was collected using the same parameters as the T1 anatomy scans.

### Cortical surface reconstructions

The cortical grey/white boundary of eight subjects was segmented automatically using FreeSurfer [37, 38] and then manually corrected using ITKGray [39]. For three subjects, segmentations were done entirely manually using ITKGray. Segmentations were installed in mrVISTA and 3D cortical surface reconstructions of the left and right hemispheres were created and displayed using mrMESH, which was also used to visualise fMRI data on the cortical surface (Stanford University, Stanford, CA; http://white.stanford.edu/software/). For Subject 4, the venous anatomy, including the Transverse Sinuses, Superior Sagittal Sinus, Straight Sinus and the confluence of sinuses, were manually segmented from the venogram using ITKGray.

### Preprocessing

Preprocessing of EPI images was performed using SPM8 (SPM software package, Wellcome Department, London, UK; http://www.fil.ion.ucl.ac.uk/spm/). Data was motion corrected using a rigid body transform and 7th degree B-spline interpolation. Images were slice scan time corrected using the first image as the reference slice and resliced into the space of the first image.

### Population receptive field (pRF) modelling

Bowtie, ring and bar stimuli were analysed using a population receptive field (pRF) method, whereby a parameterised model of the underlying neuronal response is used to calculate the predicted BOLD response. Essentially, this is a special adaptation of a linear regression, or the General Linear Model (GLM) – a standard technique in fMRI analysis [40]. The goodness-of-fit of this response is estimated using the residual sum of squares (RSS), which is minimised in a series of two-stage, coarse-to-fine searches until the optimum pRF parameters (position and size) are found. Further detail on this analysis can be found in [34].

### Fast Fourier transform and correlation analyses

A fast Fourier transform (FFT) analysis was performed on the bowties, rings and full field stimuli using inbuilt mrVISTA functions. The full field stimulus was additionally analysed using a correlation analysis between the empirical waveform and a reference waveform, which was created by convolving the stimulus timing with a canonical model of the haemodynamic response function (HRF) using AFNI’s 3dDeconvolve [41]. MATLAB was used to estimate the correlation between voxel time courses and the reference waveform, and voxels responding with ‘positive’ and ‘inverted’ HRFs were identified by the sign of their correlation value.

### Data integration across cortical depth

Data were restricted to the 3mm of voxels above the grey/white boundary (as defined by the segmentation), and averaged depth-wise across voxels to create the normalised mean, correlation and retinotopic maps.

### Mean maps

Depth-integrated, mean maps (which show the average EPI signal amplitude or EPI brightness) were constructed by voxel-wise averaging of the EPI time course and normalised by dividing the value of each voxel by the value of the voxel with the maximum intensity value. The normalised mean maps were then used to identify venous eclipses, which have a lower intensity due to the higher concentration of dHb in veins.

Regions of interest (ROIs) were hand drawn around any venous eclipse that was identified and subsequently projected onto the surface displaying the correlation analysis data to check whether voxels with inverted responses clustered in these regions. In determining whether a venous eclipse was present, we considered the shape and location of intensity drops — as venous eclipses are known to correspond with the anatomical locations of the Superior Sagittal Sinus and Transverse Sinuses, and resemble the elongated shadow of a blood vessel.

### Defining visual areas V1, V2, V3 and hV4

All visual areas were defined on the mrVista surface meshes using depth-integrated polar angle retinotopic maps as measured by the pRF modelling, with reference to the retinotopic maps generated by the FFT analysis. These two sets of maps matched up extremely well for V1, V2 and V3. The retinotopic map of hV4 was defined by a phase reversal in the polar angle maps at the lower vertical meridian or close to it, if it was not present, following the ventral hemifield model [20, 25, 28]. The ‘inverted U’ in polar angle maps demarcating the hV4/VO1 boundary, expected eccentricity reversals between hV4 and VO1 in eccentricity maps, and the location of the ptCos sulcus were also used in guiding the definition of hV4 boundaries [28, 42]. To determine the level of visual field coverage present in each hV4 map, we used a combination of visual inspection and visual field coverage plots generated by the pRF analysis.

It is well documented that voxels in regions directly outside the outermost eccentricity of the visual stimulus exhibit a ‘Negative BOLD Response’ (NBR) [43–45]. To ensure these voxels were excluded from analyses, visual area ROIs were cropped at the periphery of the mapped visual field, prior to the occurrence of NBRs. It is important here to discriminate NBRs, which typically signify a reduction in the neural response just outside the stimulated region of the visual field [43–45] from inverted voxels, which exhibit a negative correlation to visual stimulation in regions where the expected response is positive [31].

### Inverting time courses and inverted voxel counts

Once voxels with a negative correlation to the full field stimulus had been identified, custom MATLAB scripts were used to flip the time courses of these voxels in the bowtie, ring, bar and full field scans. This procedure flips the time courses of all voxels exhibiting a negative correlation, i.e. NBRs and inverted voxels. Voxels with a positive correlation were left untouched. In determining inverted voxel counts, however, it is important to note that only negatively correlated voxels within the cropped visual area ROIs were included, in order to prevent the presence of NBRs from influencing the count. Corrected time courses were reanalysed using the pRF method, resulting in a set of original and corrected pRF models. Statistical analyses on inverted voxels were conducted using SPSS version 22 [46].

### Depth-dependent analysis

To investigate whether surface vessels have a visible effect on retinotopic maps at different cortical depths, for two subjects, additional surfaces were generated at three depths using Caret v5.65 [47]. The deepest of these was generated at the grey/white boundary and expanded surfaces were created at depths of 1mm and 2.5mm from the grey/white boundary toward the cortical surface. As human visual cortex has an average thickness of 2-3mm [48], the 1mm surface is located in the mid-depth of grey matter, whilst the 2.5mm surface is close to the grey matter surface, where the large draining veins reside. These surfaces were imported into the mrVISTA session using custom written MATLAB scripts, after which they were aligned to the anatomical data using mrVista. Average time courses for the bowties, rings and full field scans were linearly interpolated from the mrVista volume space onto the depth-dependent surfaces using MATLAB. FFT analyses were performed on each interpolated time course, and correlation analyses were computed for full field time courses. Normalised mean maps of the full field data were also generated. Results were displayed on the surfaces using mrMesh. ROIs of V1, hV4 and venous eclipses were defined separately for this data, with reference to the depth-integrated retinotopic maps.

## Results

We aimed to evaluate 1) The consistency of the hypotheses that venous artefact obscures part of the lower quarterfield representation of the visual field in hV4, by resulting in either delayed [15] or inverted responses [31]; and 2) Whether it is possible to correct venous artefact in this area to more reliably measure visual field responses along the lower hV4 boundary. Our main findings were that venous eclipses did not always correspond with hV4 map coverage; the absence of venous artefact coincided with incomplete hV4 maps, whilst complete maps were found despite venous artefact being present. Furthermore, correcting inverted voxels restored a full hemifield map of hV4 in one hemisphere, but yielded only minor or no improvement in other instances of incomplete hV4 maps.

### Venous eclipses and hV4 visual field coverage

We identified visual field maps of hV4 in both cerebral hemispheres of all 11 subjects. In total, there were 8 complete and 14 incomplete hV4 maps. The left hemisphere was disproportionately represented, with all hV4 maps being incomplete, whilst only 3 right hemisphere maps were incomplete.

Using the normalised mean maps, we identified venous eclipses in 20 hemispheres, with 11 of these being on the ventral surface close to the measured hV4 maps. Of these 11, 6 were in the left hemisphere. There was no difference between the left and right hemispheres in terms of the total number of venous eclipses present.

As it has not been clearly established in the literature the extent to which artefact from the venous eclipse can spread, or the nature of the impact it may have on nearby voxels, we took a conservative approach toward classifying venous eclipses as having the ability to impact the lower hV4 boundary. As such, we considered the venous eclipse to have the ability to impact the visual field coverage of hV4 if it was present close to the hV4 map (see Supplementary materials for all normalised mean intensity and polar angle maps). Despite this conservative approach, we did not find a clear relationship between the presence of a venous eclipse near hV4 and the degree of visual field coverage represented. Whilst incomplete maps of hV4 did correspond with the presence of a venous eclipse in some hemispheres, we identified instances where an incomplete hV4 map was identified despite the absence of a venous eclipse, and likewise instances of complete maps of hV4 in the presence of a venous eclipse. These results are summarised in Fig 3.

**Fig 3.**
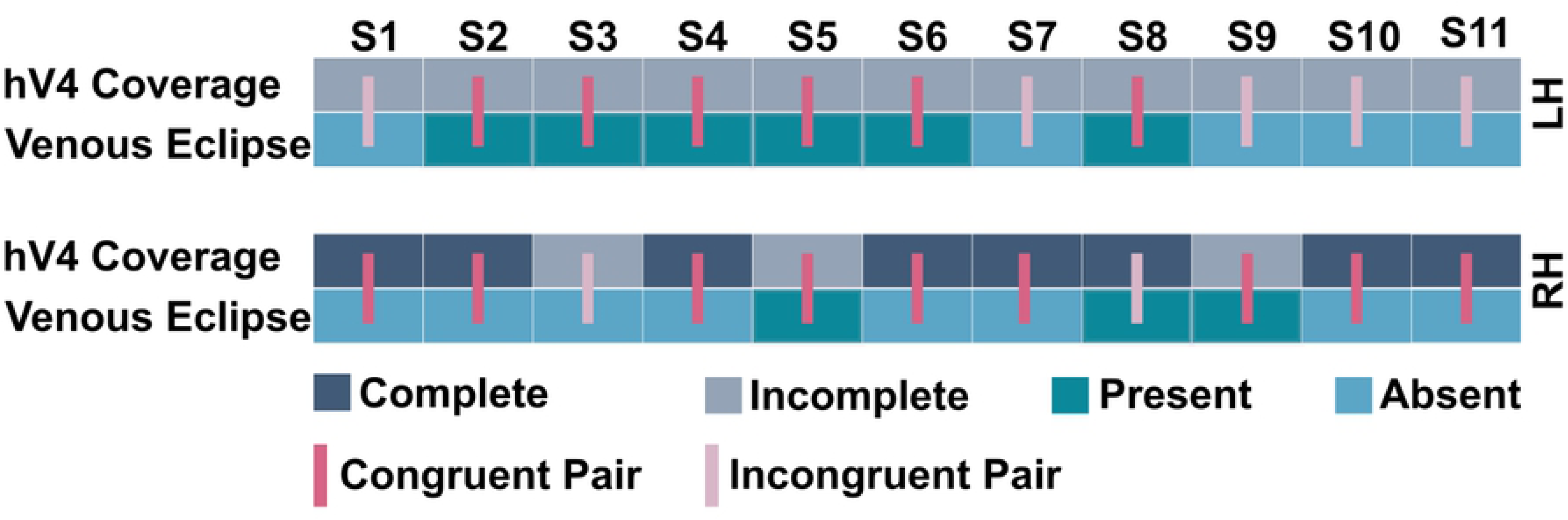
Intersection of hV4 map coverage and venous eclipse presence. Visual field coverage is classified as ‘complete’ when the representation of the visual field continued to the lower vertical meridian (or close to it), and ‘incomplete’ when it only extended to approximately three quarters of the contralateral visual hemifield or less. Venous eclipses were considered present if they were located proximate to the defined hV4 map, and absent if further away or non-existent. Congruent pairings were those with a coincidence of *incomplete hV4/Venous eclipse present* or *complete hV4/Venous eclipse absent*. Incongruent pairings were those with a coincidence of *complete hV4/Venous eclipse present* or *incomplete hV4/Venous eclipse absent*.

### Inverted voxels

Inverted voxels were present in all visual areas examined, and always (though not exclusively) clustered in regions corresponding with venous artefact (Fig 4). It is important to note that inverted voxels did not appear to strictly adhere within the region of the venous eclipse, as seen in the normalised mean maps, with inverted voxel clusters extending into neighbouring regions.

**Fig 4.**
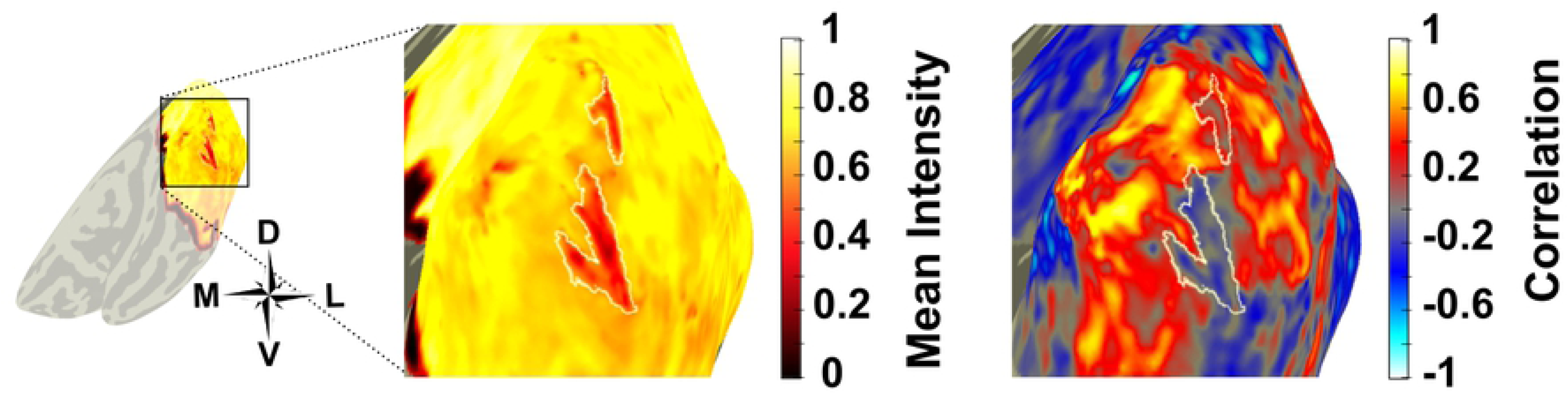
Cluster of inverted voxels in the region of the venous eclipse.

Inverted voxels tended to cluster in regions of the venous eclipse, as shown here in the right hemisphere of Subject 5. The insert to the left shows the entire cortical surface; the highlighted section indicates the magnified area to the right. A) The venous eclipse ROI is defined in white on the normalised mean map; B) The correlation map shows a cluster of voxels with a negative correlation represented in blue. The representation of inverted voxels extends beyond the region the of venous eclipse as it appears in the mean map, however it should be noted that the region in the mean map corresponding to the inverted voxels still shows a lower mean intensity value than regions corresponding to positive voxels.

A 2×4 repeated measures ANOVA was used to test for differences in the percentage of inverted voxels between the left and right hemispheres and each visual area. A significant main effect of visual area was found (F3,30 = 4.84, p = .007), with follow up analyses showing inverted voxel percentages in V1 were significantly lower than V2 (F3,30 = 7.81, p = .035) and V3 (F3,30 = 7.81 p = .005) in the right hemisphere. No other significant differences were present. We find similar percentages of inverted voxels in hV4 as Puckett and colleagues, however higher percentages in V1, V2 and V3 [31]. Percentages of inverted voxels across visual areas and hemispheres can be seen in Table 1.

**Table 1.** Normalised percentages of inverted voxels in visual areas V1, V2, V3 and hV4, averaged across subjects.

| Visual Area | V1 |  | V2 |  | V3 |  | hV4 |  |
| --- | --- | --- | --- | --- | --- | --- | --- | --- |
| Hemisphere | LH | RH | LH | RH | LH | RH | LH | RH |
| Total#Voxels | 42 078 | 36 723 | 35 871 | 38 027 | 31 034 | 34 212 | 11 588 | 10 124 |
| %Inverted | 6.74 | 6.65 | 10.91 | 12.88 | 9.02 | 11.44 | 10.60 | 7.94 |

Impulse response functions (IRFs) were calculated for the full field stimulus; no differences between the left and right hemisphere IRFs were apparent, therefore data displayed here is collapsed across hemispheres. Fig 5 shows that, in contrast to Puckett and colleagues [31], voxels exhibiting an inverted BOLD response in hV4 have the lowest percent signal change of all areas. A lower percent signal change is also present in V1-V3 compared to positive voxels. Positive responses did not vary across visual areas, with positive voxels exhibiting a consistent time-to-peak of 8s after stimulus onset in all visual areas. Inverted responses showed some slight variation, with a time-to-peak of 6s for V1-V3, while the time-to-peak for V4 was 8s.

**Fig 5.**
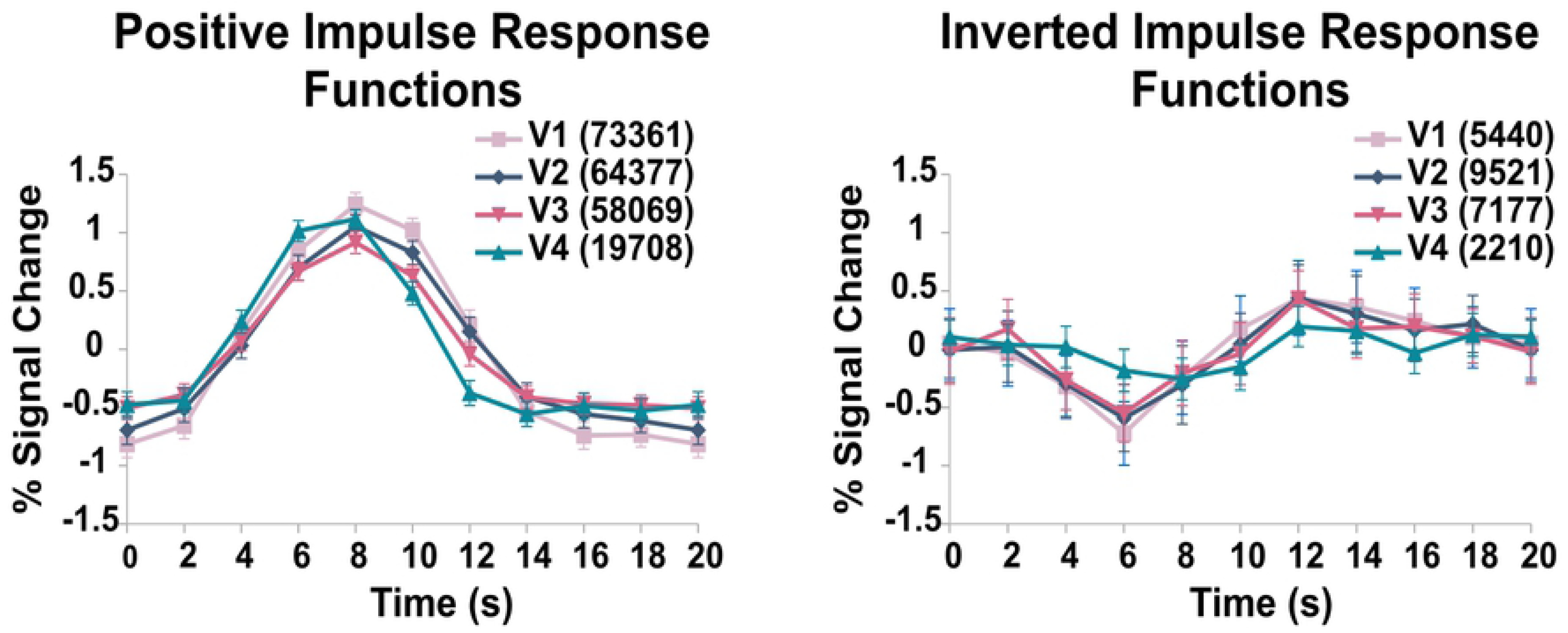
Positive and inverted impulse response functions. Average single cycle full field responses across all subjects and visual areas, collapsed across hemispheres. Voxel counts for each visual area are displayed in brackets on the legend. Error bars represent +-1 SEM.

### Correcting retinotopic maps

Correcting inverted voxels did not have a major effect on the number of complete hV4 maps, however in one subject there was a restoration of a complete ventral hemifield following correction (Fig 6). Small changes were seen in the hV4 maps of all other subjects, most of which were inconsequential, however minor improvements were present in some (Supplementary Figure 1).

**Fig 6.**
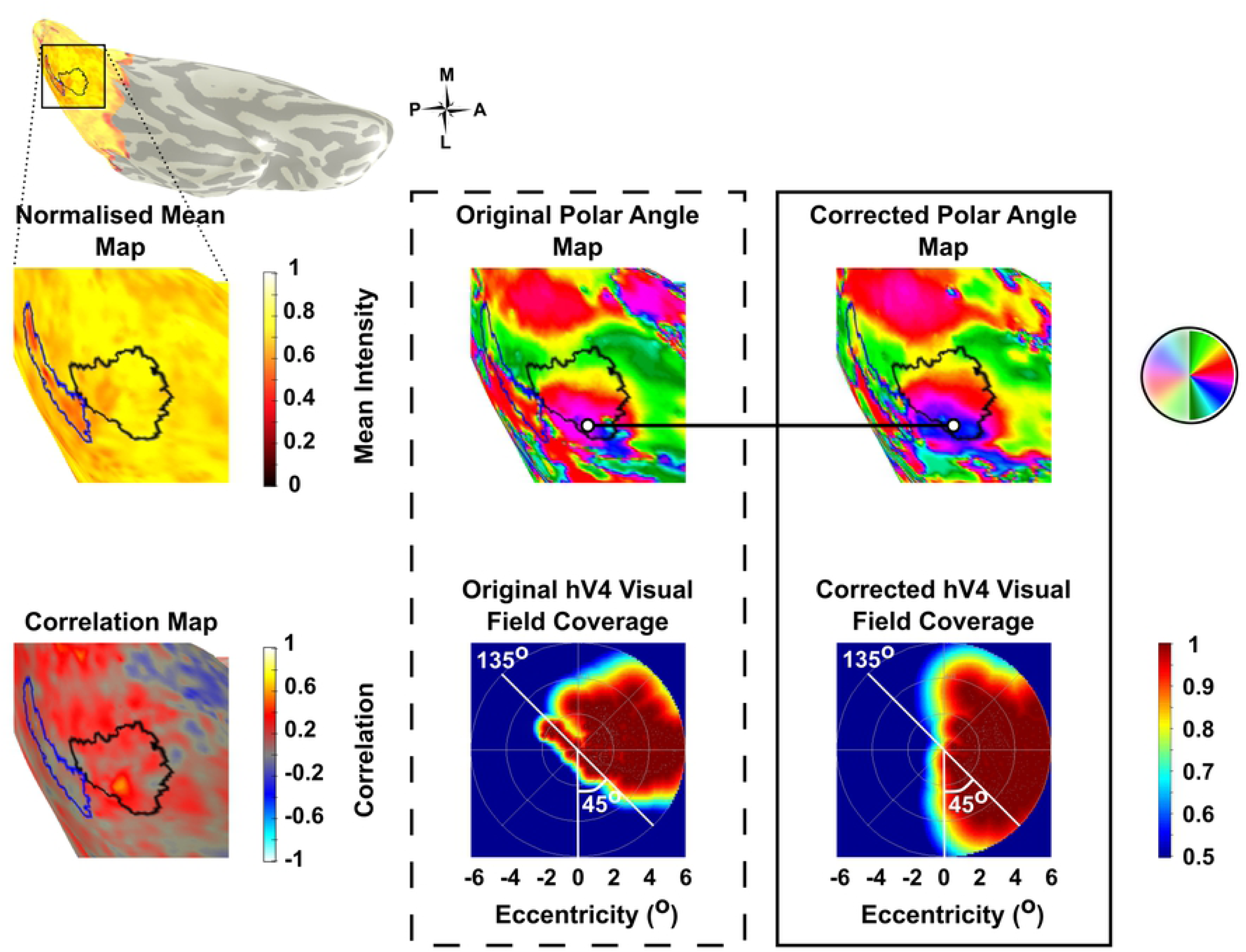
Restoration of hV4 in Subject 10 after correcting inverted voxels. The cortical surface of Subject 10 (left hemisphere, viewed ventrally) is inserted in the upper left. Mean intensity, correlation and polar angle maps show venous eclipse (blue) and hV4 (black) ROIs. Venous artefact in the mean map is discernible by regions of lowered intensity. The venous artefact in the correlation map is characterised by low and negative correlations with model HRFs. The middle and right columns show results from a pRF analysis, with polar angle maps on top and visual field coverage plots from the hV4 ROI below. Note the changes in the pRF analysis before (dashed line) and after (solid line) correcting inverted voxels (white dots). Original pRF data shows the polar angle representation of hV4, which is missing approximately the lower 45°, reflected in the corresponding visual field coverage plot, where circles represent individual pRFs. In the corrected pRF map, hV4 coverage extends further, almost reaching the lower vertical meridian, which is reflected in the visual field coverage plot. Correcting inverted voxels has resulted in a more accurate map of hV4, based on visual inspection and visual field coverage. Note the incorrect coverage of the ipsilateral hemifield at 135° present in the original map is no longer a problem in the corrected map. Most significantly, the visual field coverage plot of the corrected map has a representation of the full contralateral visual hemifield.

Although our main hypotheses concerned hV4, an ancillary effect of performing the correction procedure on negatively correlated voxels outside this area was noted for early visual areas (V1, V2 and V3). Instead of corresponding to the venous eclipse, a large number of voxels exhibiting a negatively correlated response to full field stimuli could be classified as NBRs. These represent neuronal suppression, and are usually located just outside (and in some cases just within) the stimulated region [44, 45, 49].

Across V1-V3 ROIs, we found NBRs located just outside the eccentricity range of visual stimulation (i.e. less than 5.5°eccentricity). These voxels corresponded to correlation values that were almost as strong in the negative direction as positively correlated voxels that were within the range of visual stimulation. Large improvements in phase maps were observed at the periphery of V1-V3 ROIs when flipping the time courses of NBRs. This was particularly notable in V1, where large clusters of NBRs were present in regions immediately adjacent to the region of cortex being stimulated. The correction (i.e. the flipping of NBR time courses) had a striking effect, where coverage of the visual field map was extended beyond the furthest eccentricity of the stimulus. This correction procedure demonstrates the potential to retinotopically map visual cortex beyond the often limited eccentricity range of MR visual display systems (Fig 7).

**Fig 7.**
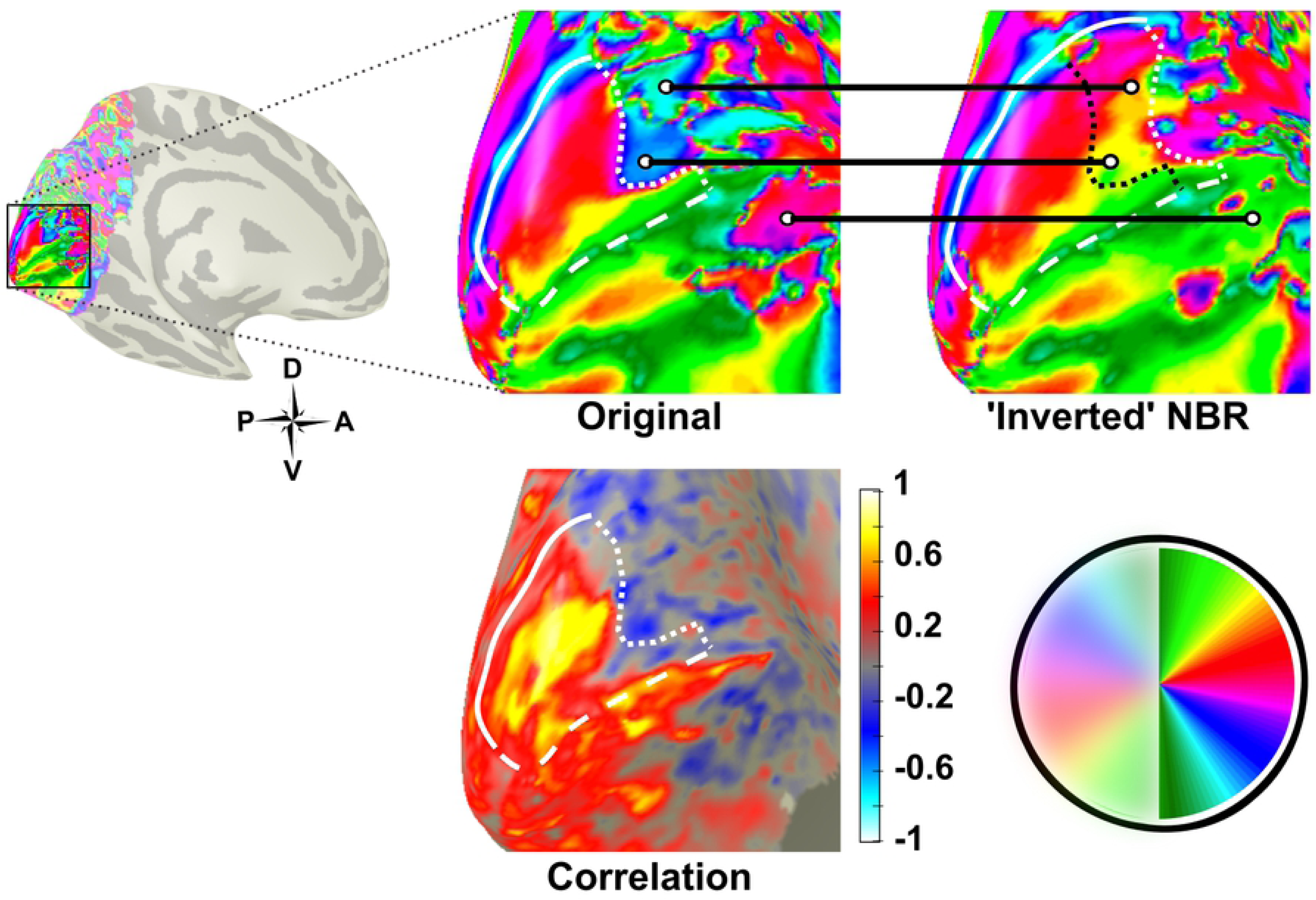
Retinotopic map of V1 in the left hemisphere of Subject 4 before and after flipping negatively correlated time courses. Insert on the left shows the entire cortical surface; highlighted section indicates magnified views to the right. The original pRF map is displayed with the V1 ROI in white. Regions where the order of the visual field representation is ‘inverted’ are indicated by white circles, which attach to the same regions in the ‘corrected’ V1 map. Here it can be seen that significant extensions to the V1 map are present after flipping NBR time courses. Correlations of NBRs are shown in the correlation map.

### Depth-dependent effects of venous artefact on retinotopic maps

To extend our analysis of the effect of venous artefact on retinotopic maps, we performed a post hoc investigation of its impact across cortical depth (Fig 8). As such, for two subjects we made cortical surfaces at three depths (grey/white, 1mm and 2.5mm above the grey/white boundary). Normalised mean maps for all 2.5mm surfaces show clearly, and in detail, what appear to be venous eclipses caused by the SSS and TSs. This is in contrast to the depth-integrated mean maps (shown in Figures 4 and 6), where averaging data across 3mm of grey matter seems to have the effect of weakening the appearance of the venous eclipse, to the extent that it disappears entirely in some regions. The venous eclipses increasingly weaken in appearance with depth, which can be clearly seen in all of the 1mm and grey/white surfaces.

**Fig 8.**
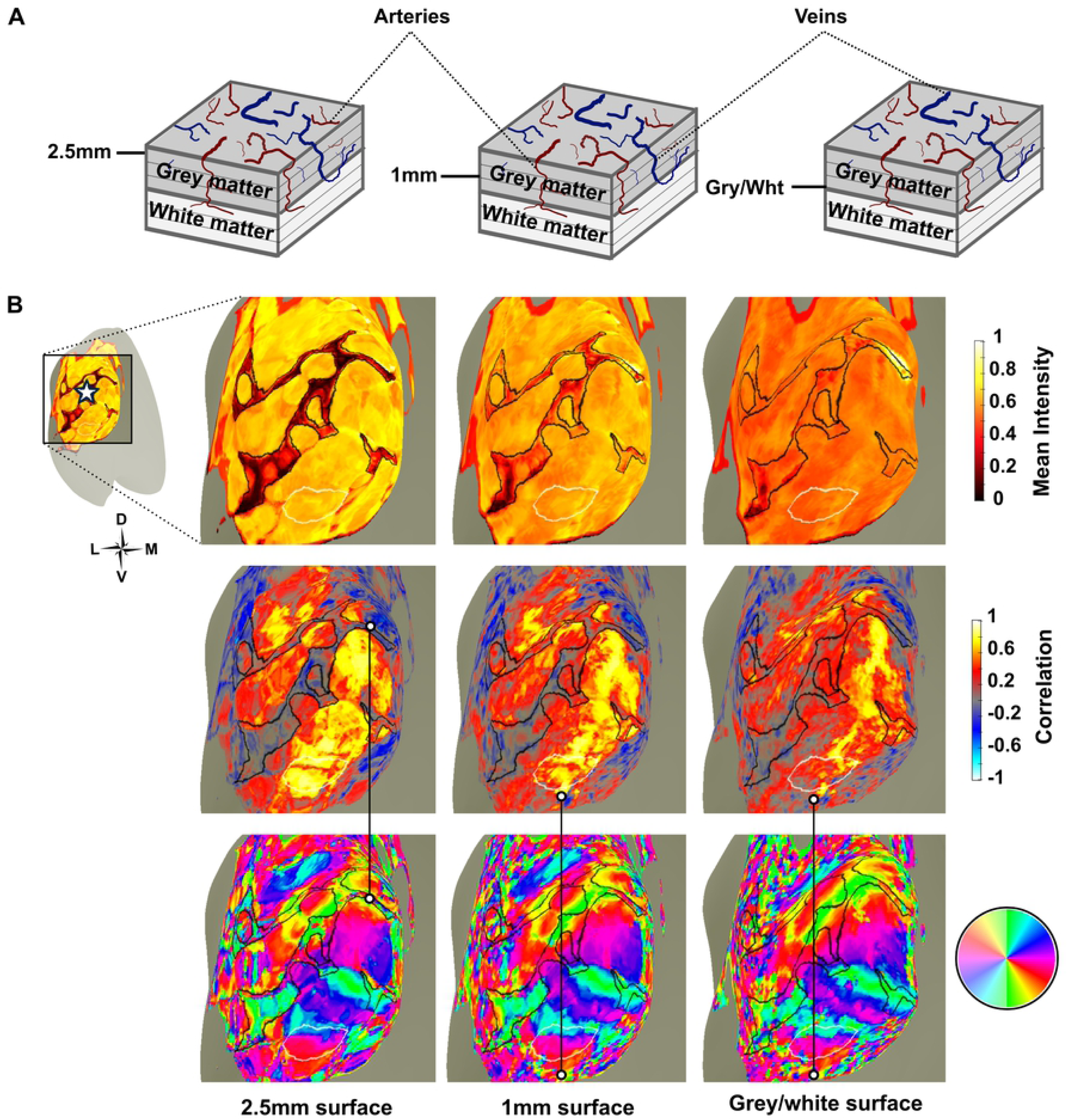
Depth-dependent effects of the venous eclipse in the left hemisphere of Subject 6. A) Schematic of layers of the cerebral cortex depicting cortical arteries (red) and veins (blue). These can be seen diving through the layers of grey matter before turning at 90° upon reaching the white matter and thinning into branches [51]. B) Depth-dependent surfaces at 2.5mm, 1mm and the grey/white boundary showing the venous eclipse (black ROI) and hV4 (white ROI). The star in the top left insert represents the occipital pole. Mean intensity maps (top) show the venous eclipse weakening with grey matter depth, and a corresponding reduction in the number of inverted voxels within the venous eclipse ROI is seen in the correlation maps (middle; highlighted by white dots in Column 1). The polar angle maps (bottom) show an incomplete map of hV4, which does not change across depth (Note, the ‘speckles’ indicated by the white dots in the second and third columns highlight what could be the lower vertical meridian of the hV4 map, however as the correlation map shows multiple inverted voxels in this region, it is difficult to trust the responses of these voxels.

A reduction in the mean intensity of all voxels is apparent with decreasing cortical depth. This reflects the fact that the BOLD signal is generally stronger in layers of grey matter closer to the pial surface [50]. Strikingly, voxels in regions surrounding the venous eclipse show a high mean intensity despite being immediately adjacent to venous artefact. This is particularly evident at the more superficial depths.

Compared to the depth-integrated data, inverted voxels are more tightly restricted within the venous eclipse, and not outside it in the 2.5mm surfaces. This indicates that whilst voxels immediately beneath surface vessels respond with abnormal time courses, voxels immediately adjacent to, but outside this region have normal responses to full field stimulation. This is also reflected in the mean intensity of these voxels, which is comparable to those which are far away from venous artefact (see Fig 8B). Furthermore, the clusters of inverted voxels within the venous eclipse ROI decrease in frequency with grey matter depth. Other, more sparsely distributed voxels exhibiting negative correlations appear in regions distant from the venous eclipse in the 1mm and grey/white surfaces, increasing in frequency with depth (Fig 9).

**Fig 9.**
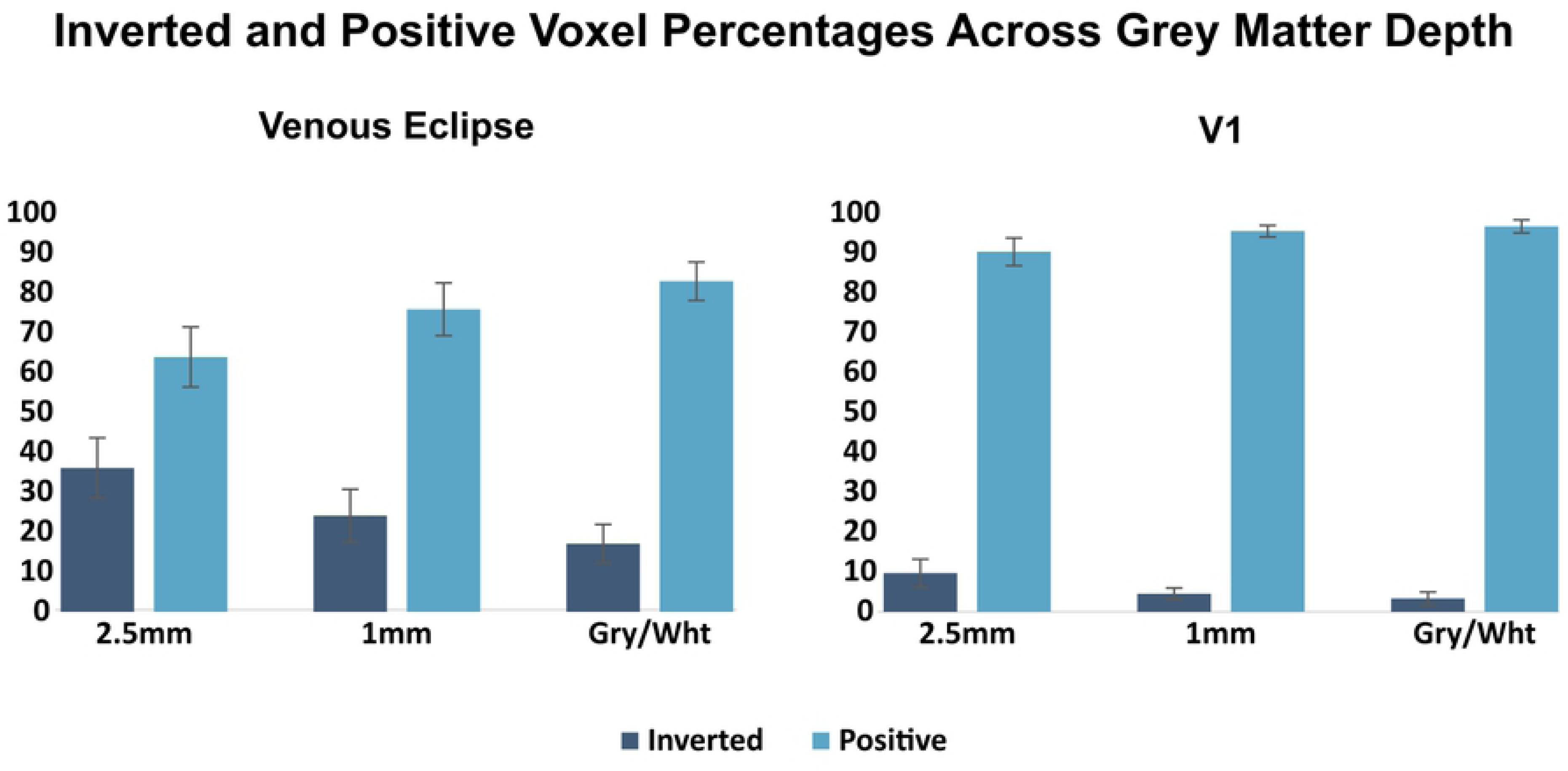
Average percentage of inverted and positive voxels in the venous eclipse and V1 ROIs for the left and right hemispheres of Subjects 5 and 6. A reduction in the mean percentages of inverted voxels within the venous eclipse and V1 is apparent as one progresses from the cortical surface toward the Grey/White boundary, with the venous eclipse having higher percentages of inverted voxels in all surfaces compared to V1. Error bars represent +-1 SD.

## Discussion

Our primary aim was to evaluate the consistency of the venous eclipse and inverted voxels as limitations to accurately measuring retinotopic maps of hV4 [15, 31]. We report results pertaining to hV4 maps, venous eclipses and inverted voxels, as examined in depth-integrated and depth-dependent data. We also illustrate the utility of surface maps that are localised at specific grey matter depths and suggest that such maps are a useful tool in studying or avoiding the disturbances that blood vessels can cause to MRI analysis.

### Correcting inverted voxels can restore the lower boundary of hV4 in some hemispheres

Consistent with previous studies [20, 22, 24, 27, 28], we found that receptive field centres in hV4 maps often do not extend over the entire hemifield, with more than half (64%) of hV4 maps being incomplete. Attempts to restore these maps by flipping the time courses of inverted voxels showed that it is possible to improve some of the incomplete maps, even restoring an entire hemifield in one instance. This singular case serves as a clear example of a subject who has a complete hemifield representation in hV4 which was obscured by an inadequacy of standard BOLD fMRI measurements.

Our successful correction of this hV4 map highlights that some incomplete hV4 maps seem to be caused by inverted voxels being present along the lower boundary. These inverted voxels may be the result of venous artefact, and can be accounted for using appropriate techniques. However, we were unable to restore the lower boundary of most incomplete hV4 maps, despite improving the order (i.e. smoothness) of the visual field representation in some (see Supplementary Fig 1).

### Venous eclipses do not always coincide with incomplete hV4 maps

We found little concordance between hV4 map coverage and the presence of venous eclipses in our depth-integrated mean intensity maps, with seven incongruent pairings existing in our data. Furthermore, we find six cases of incomplete hV4 maps with no observable venous eclipse in the vicinity (LH of S1, S7, S9, S10, S11 and RH of S3), close to half of the incomplete maps we identified.

A potential explanation for this unexpected discrepancy is that the intensity drop characteristic of venous artefact may be limited to voxels in the upper layers of grey matter, due to their closer proximity to surface veins. Averaging the responses of these superficial voxels with those from deeper layers of grey matter may obscure or weaken the amplitude drop of the venous eclipse to the extent it disappears or becomes difficult to identify. Essentially, not seeing venous artefact clearly in mean intensity maps may not signify its absence. Our depth-dependent surfaces support this idea, as they show the venous eclipse fading from being clearly recognisable in the 2.5mm surface to weakening in some parts of the 1mm surface, and disappearing entirely in most sections of the grey/white surface.

### Clusters of inverted voxels are present in the region of the venous eclipse

We found inverted voxels to cluster in and around the venous eclipse in most hemispheres (see Fig 4), replicating the same finding by Puckett and colleagues [31]. While it seems clear that inverted voxels are present at higher frequencies near venous artefact, it is difficult to ascertain whether these voxels are located outside the venous eclipse or merely in interstices created by the averaging of data from multiple grey layers (see above). It is possible that these inverted voxels are in fact contaminated by venous artefact, which is unidentifiable in the depth-integrated mean intensity maps. An alternative possibility is that the venous eclipse spuriously affects the time courses of voxels that fall laterally to its location, in addition to beneath it.

Our depth-dependent results suggest that clusters of inverted voxels are reasonably restricted to the regions of grey matter which show venous artefact, especially in the 2.5mm laminar surface. Responses directly neighbouring the outside of the venous eclipse in this layer show both high mean intensities and strong positive correlations, indicating that voxels which do not form part of the venous eclipse are not impacted by it. However more depth-dependent analyses are required to confirm and better quantify this finding.

### Inverted voxels affect most visual areas equally

In contrast to previous work [31], we do not find percentages of inverted voxels to be higher in V4 compared to V1, V2 and V3. We additionally do not find inverted voxel percentages to differ between the left and right hemisphere; this is consistent with absence of differences in the number of venous eclipses between hemispheres. This discrepancy with earlier work may be due to our larger pool of young adult subjects (n=11, aged 21-26), compared to Puckett and colleagues, whose sample differed in size and age range (n=4, aged 21-61).

This is a potentially important difference, as neurovascular decoupling (where neural activity is not accompanied by compensatory changes in the dilation and constriction of blood vessels) has been reported to induce inverted BOLD responses in regions where blood vessels are impeded as a function of ageing and disease [52, 53]). The smaller number of subjects in Puckett and colleagues’ work may also explain this, as the TS has some natural variability in its proximity to hV4 [15]. The presence of a larger percentage of inverted voxels in one hV4 map due to the TS more proximate to it may skew the overall findings more easily than in our pool of 11 subjects.

### The pattern of inverted voxels changes with surface depth

Within the venous eclipse in our depth-dependent data, the number of inverted voxels appears to decrease with depth; many inverted voxels in the 2.5mm surface have positive correlations in the 1mm and grey/white surfaces. Interestingly, outside the venous eclipse smaller, sporadic inverted voxels appear in deeper layers where the corresponding voxels in the surfaces above show positive correlations.

We cannot say with certainty what may be causing these isolated inverted voxels, however they may simply be due to a lower SNR in deeper grey matter, increasing the frequency of random negative correlations. The sparse distribution and smaller clusters of these inverted voxels may also be the result of small veins from lower cortical layers, which dive through the grey matter from the cortical surface, bending at a right angle upon reaching the white matter before branching back up into lower grey layers (see Fig 8A) [51].

Regardless of the source of these inverted voxels, their impact can be clearly seen in the polar angle maps, where affected regions appear to map non-neighbouring areas of the visual field, disturbing the order of human visual cortex. This is in contrast to what is known about the organisation of this region of the human brain, and in agreement with previous work [31].

### Hemispheric asymmetry in hV4 map coverage cannot be explained by a corresponding asymmetry in venous anatomy

Having been noted in previous work and corroborated here, the bias towards left hemisphere hV4 maps being incomplete more frequently than right hemisphere maps is a robust finding that warrants explanation. Given that the TS and large surface veins in general give rise to the venous eclipse, consideration is due to the physical anatomy of veins in the occipital cortex. Individual variations exist at the torcular Herophili (tH; the confluence of sinuses at the occipital pole) [54, 55], where variations are separated into three broad categories, depicted in Fig 10. Of particular note is the Type 1 where there is an *absence* of a confluence, and the right and left transverse sinuses connect to only one of the Superior Sagittal Sinus (SSS) and Straight Sinus (SS) [54, 55] (see ‘Type 1’ in Fig 10).

**Fig 10.**
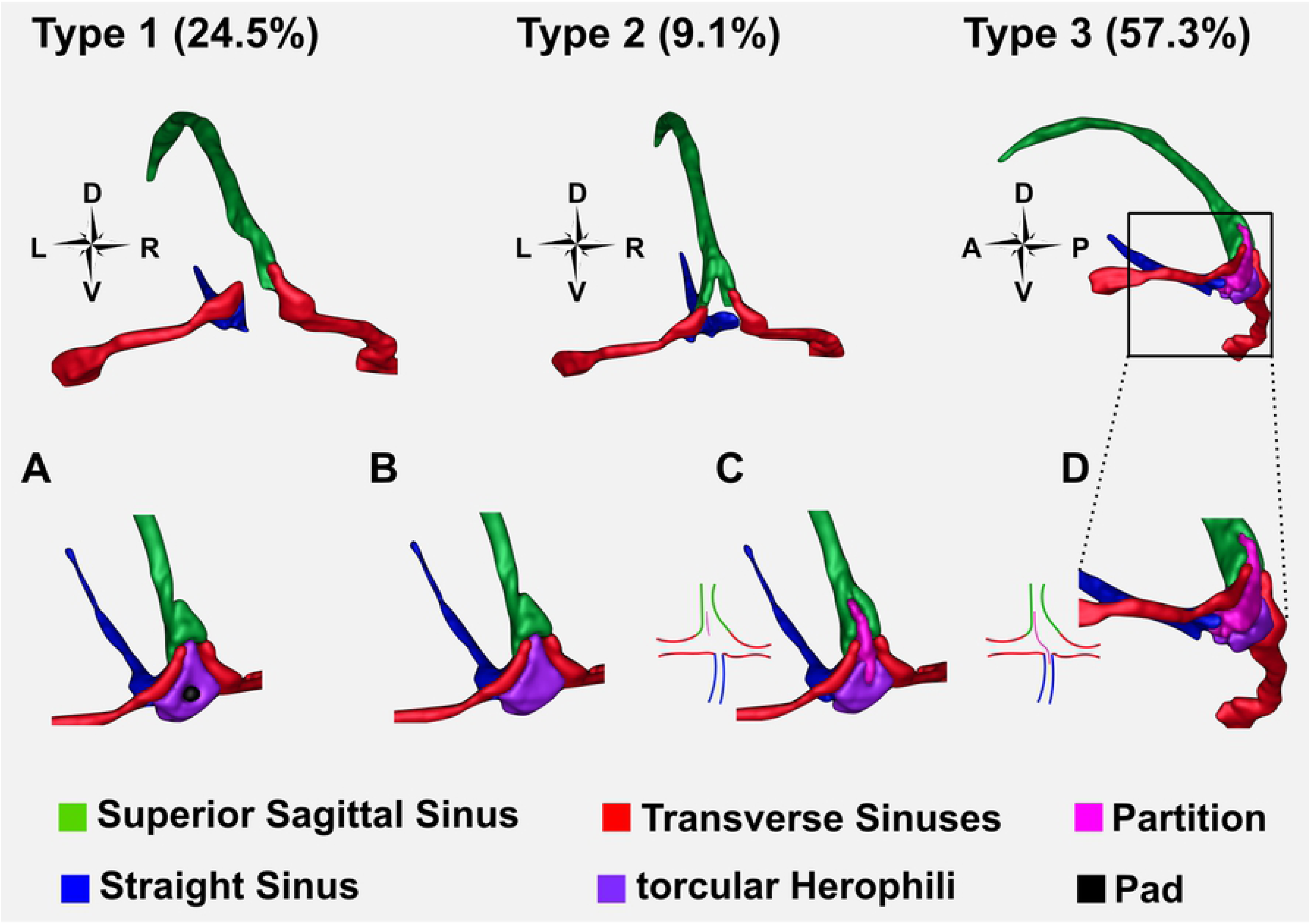
Three main variations on the venous anatomy of the dural sinuses. Percentages refer to the frequency with which each type is seen. In Type 1, the Superior Sagittal Sinus connects to one Transverse Sinus, and the Straight Sinus to the other, with the two TSs completely separate from one another. In Type 2, the SSS and SS are forked, with the left forks connecting to the left TS, and the right forks to the right TS. In Type 3 there is some variation of a confluence of the sinuses. A) A ‘pad-like dural elevation’ in the tH [54]. B) Sinuses are connected by a confluence at the occipital pole. C) A partial partition extends from the SSS into the confluence of sinuses. Partition can be closer to the left or the right wall of the SSS. D) A full partition extends from the SSS diagonally through the tH to the SS. Full partition can extend diagonally across from closer to the left or the right wall of the SSS. Figure was creating by segmenting the venous anatomy from the venogram of Subject 4, who had Type 3A. Other variations were approximated based on the descriptions in [54].

This is important to note, as research has shown the signal recorded in areas of large draining veins may be reflective of changes in distal regions of cortex [14]. As such, in subjects where the SSS and SS drain into separate TSs, the responses being measured in the region of the TS come from non-homologous regions of each cerebral hemisphere. Specifically, the SSS would be fed by lateral regions of the anterior cerebral hemisphere, and the SS by the cerebellum and centre of the head [18]. For hV4 maps belonging to subjects with this venous anatomy, signal recorded in the region of hV4 in the left and right hemispheres may be asymmetric, as the blood in the TSs would be originating from two distinct brain regions. However it is important to note that Type 1 represents only 24.5% of cases, and this considerable variability of TS anatomy is difficult to reconcile with the relatively consistent finding of incomplete left and complete right hemisphere hV4 maps.

Alternatively, a bias in TS size, whereby the left TS tends to be larger could serve as a potential explanation, as a higher concentration of deoxyhaemoglobin near left hemisphere hV4 maps should increase the chances of finding incomplete hemifields. However an investigation of 110 adult cadavers found the opposite – a (small) bias towards the right TS being larger – a finding that has been replicated recently [54, 55]. Together this suggests that a strong association between hV4 map coverage and venous anatomy is yet to be consolidated and further work in this area is required.

### Susceptibility artefacts may account for hemispheric asymmetry

As there is a low likelihood that hV4 maps vary so considerably in their representation of the visual field between the left and right hemispheres as found here and elsewhere [20, 22, 24, 27, 28, 56], venous asymmetry is one of multiple potential causes. Another possible explanation of the hemispheric asymmetry in our hV4 maps is the fact that our phase encoding direction (also known as the ‘blip’ or ‘fat-shift’ direction) was left to right.

This has the effect of distorting the EPIs in the same direction, as a result of susceptibility artefacts induced by various types of tissue [57, 58]. However, though sufficient as a hypothesis for our data, the same leftwards asymmetry reported by Winawer and colleagues cannot be attributed to a left to right phase encoding direction, as this data was acquired using a spiral sequence [15]. Nevertheless, the direction of phase encoding is worth taking into consideration when interpreting findings of hemispheric asymmetry in fMRI data, particularly when no known explanation for the asymmetry exists.

### Negative BOLD responses versus inverted voxels

Thus far, we have been considering the impact of inverted voxels on retinotopic maps; it is important to distinguish these inverted voxels from Negative BOLD Responses (NBRs). While many details of the specific drivers of NBRs are still a matter of debate, it is likely that they are due to neural mechanisms and a reduction of neuronal activity [31, 44, 45, 59]. Despite having causal differences, both NBRs and inverted voxels show a negative correlation with Full Field visual stimuli; however, we observe a range of differences between voxels showing NBRs and voxels with ‘inverted’ responses.

In our study, voxels showing NBRs are arranged in large clusters located adjacent to and outside the region of cortex that fell within the eccentricity of our stimuli. They have stronger responses than inverted voxels, often as strong as adjacent PBRs. This strong and reliable response enabled the convincing extension of retinotopic maps of early visual areas, after flipping negatively correlated time courses, which allowed for the delineation of the visual field map of V1 beyond the range of our visual display system.

### Limitations/Considerations from the depth-dependent data

Examination of the depth-dependent data shows that the venous eclipse is most prominent in superficial surfaces, which is to be expected as these are in close proximity to surface veins. Based on this depth-dependent analysis, it appears that the grey/white boundary may be impacted very little by venous artefact, at least in terms of the mean intensity of voxels in cortex underlying surface veins. The polar angle maps depicted in Fig 8 clearly show an incomplete map of hV4 at the superficial surface, despite the fact there is no venous artefact visible along the lower boundary, and where mean intensity and correlation responses are comparable to surrounding regions that are unaffected by venous artefact. At the 1mm and grey/white depths, the map of hV4 still appears to be incomplete.

Careful examination of the region surrounding the lower hV4 boundary shows some voxels which appear to respond to the lower vertical meridian, however as the correlation maps show that these voxels also exhibit weak or inverted responses, it is difficult to trust the retinotopic mapping in this region. It must be noted that the conservative criteria we used in classifying whether hV4 maps were likely to be impacted by venous artefact resulted in the inclusion of this data as a congruent *Incomplete hV4/Venous eclipse present* pairing, based on depth-integrated maps. Examination of this data across three different grey matter depths indicates that the venous eclipse could not reasonably be expected to impact responses along the lower hV4 boundary, and hence this hemisphere might be more accurately classified as an incongruent *Incomplete hV4/Venous eclipse absent* pairing.

Although the depth-dependent analysis formed only a peripheral part of this work, we believe that our findings here are important for several reasons. Firstly, we show that depth-dependent analyses can be performed on medium resolution fMRI data (1.5mm), and that doing so yields additional, meaningful information about the responses of voxels at different cortical depths. Most importantly we demonstrate that it is possible to locate regions that are impacted by what appears to be a venous eclipse with a high level of spatial specificity. In future work, this will enable more accurate predictions of where venous artefact can be expected to impact retinotopic maps in other fMRI analyses, including beyond the visual cortex. Finally, our results suggest that regions outside the venous eclipse do not appear to be impacted by it, and deeper cortical layers show only minor impacts of surface veins on voxel responses.

## Conclusion

We demonstrate a hemispheric asymmetry that is biased towards incomplete maps of hV4 appearing predominantly in the left hemisphere, a finding that has never been thoroughly discussed despite being corroborated in multiple earlier studies, and for which an explanation is wanting [20, 22, 24, 27, 28]. The leftwards bias found for incomplete hV4 maps cannot be explained by a corresponding anatomical bias of the TS [54], suggesting that venous eclipses can explain some, not all, incomplete hV4 maps. It appears to be the case that an additional, more consistent source of the hemispheric asymmetry must exist to explain these incomplete maps, and we suggest this cause should be more consistently biased towards the left.

Venous eclipses and inverted voxels have both been proposed as explanations for incomplete maps of hV4 [15, 31]. We confirm and support previous findings identifying cases where inverted voxels cluster in regions of the venous eclipse, and we show that correcting time courses of inverted voxels restored a complete hemifield map of hV4 on the ventral surface in one hemisphere as well as improved the disrupted retinotopic maps in others. This strongly supports the notion that hV4 maps are complete but may appear incomplete due to inadequacies in measurement.

## Acknowledgements

We thank Professor Robert Barry for helpful suggestions. This research has been conducted with the support of an Australian Government Research Training Program Scholarship.

## Supplementary Materials

**Supplementary Fig 1. Changes in hV4 maps after flipping the time courses of negatively correlated voxels.** The inserts on the left show the entire cortical surface; square indicates magnified views to the right. A) Left hemisphere maps from Subject 6. B) Left hemisphere maps from Subject 8. Black dots indicate regions of improvements.

**Below are the mean intensity, correlation, hV4 polar angle maps and pRF coverage plots from all subjects. Asterix indicates subjects with only one repetition of each stimulus (25 minutes scan time).**

## References

1. Felleman DJ. Visual System in the Brain. International Encyclopedia of the Social & Behavioral Sciences. 2001; p. 16278–16285. doi:10.1016/B0-08-043076-7/03473-2.

2. Grill-Spector K, Malach R. The human visual cortex. Annual Review of Neuroscience. 2004;27(1):649–677. doi:10.1146/annurev.neuro.27.070203.144220.

3. Gulyas B, Ottoson D, Roland PE. Functional Organisation of the Human Visual Cortex. vol. 61 of Wenner–Gren International Series. Pergamon; 1993.

4. Zeki S. Functional specialisation in the visual cortex of the rhesus monkey. Nature. 1978;274:423–428.

5. Engel SA, Glover GH, Wandell BA. Retinotopic organization in human visual cortex and the spatial precision of functional MRI. Cerebral cortex. 1997;7(2):181–192.

6. Holmes G. Disturbances of vision by cerebral lesions. British Journal of Ophthalmology. 1918;2(7):353–384.

7. Inouye T. Die Sehstorungen bei Schußverletzungen der kortikalen Sehsphäre: nach Beobachtungen an Verwundeten der letzten japanischen Kriege. W. Engelmann; 1909.

8. Barlow HB. Why have multiple cortical areas? Vision Research. 1986; p. 81–90.

9. Schira MM, Tyler CW, Rosa MGP. Brain Mapping: The (Un)Folding of Striate Cortex. Current Biology. 2012;22(24):R1051 – R1053. doi:https://doi.org/10.1016/j.cub.2012.11.003.

10. Wandell BA, Winawer J. Imaging retinotopic maps in the human brain. Vision Research. 2010;51:718–737.

11. Boubela R, Kalcher K, Huf W, Seidel EM, Derntl B, Pezawas L, et al. FMRI measurements of amygdala activation are confounded by stimulus correlated signal fluctuation in nearby veins draining distant brain regions. Nature. 2015;5:10499–10515.

12. Curtis AT, Hutchison RM, Menon RS. Phase based venous suppression in resting-state BOLD GE-fMRI. NeuroImage. 2014;100(Supplement C):51–59. doi:https://doi.org/10.1016/j.neuroimage.2014.05.079.

13. Menon RS. The great brain versus vein debate. NeuroImage. 2011;62(2):970–974. doi:10.1016/j.neuroimage.2011.09.005.

14. Olman C, Inati S, Heeger D. The effect of large veins on spatial localization with GE BOLD at 3 T: Displacement, not blurring. NeuroImage. 2007;34:1126–1135.

15. Winawer J, Horiguchi H, Sayres RA, Amano K, Wandell BA. Mapping hV4 and ventral occipital cortex: The venous eclipse. Journal of Vision. 2010;10:1–22.

16. Duyn J, Frank J, Ramsey N, Mattay V, Sexton R, Tallent K, et al. Effects of large vessels in functional magnetic resonance imaging at 1.5T. International Journal of Imaging Systems and Technology. 1995;6(2-3):245–252. doi:10.1002/ima.1850060216.

17. Ogawa S, Lee T. Magnetic Resonance Imaging of Blood Vessels at High Fields: In Vivo and In Vitro Measurements and Image Simulation. Magn Reson Med. 1990;16:9–18.

18. Gray H, Lewis WH. Anatomy of the human body. Philadelphia, Lea & Febiger; 1918.

19. Arcaro MJ, McMains SA, Singer BD, Kastner S. Retinotopic Organization of Human Ventral Visual Cortex. Journal of Neuroscience. 2009;29(34):10638–10652. doi:10.1523/JNEUROSCI.2807-09.2009.

20. Brewer A, Liu J, Wade A, Wandell B. Visual field maps and stimulus selectivity in human ventral occipital cortex. Nature Neuroscience. 2005;8(8):1102–9. doi:10.1038/nn1507.

21. Hadjikhani N, Liu AK, Anders D, Cavanagh P, Tootell R. Retinotopy and color sensitivity in human visual cortical area V8. Nature neuroscience. 1998;1(3):235–241.

22. Hansen K, Kay K, Gallant J. Topographic Organization in and near Human Visual Area V4. The Journal of Neuroscience. 2007;27:11896–11911.

23. McKeefry D, Zeki S. The position and topography of the human colour centre as revealed by functional magnetic resonance imaging. Brain. 1997;120(12):2229–2242.

24. Tootell R, Hadjikhani N. Where is ‘Dorsal V4’ in Human Visual Cortex? Retinotopic, Topographic and Functional Evidence. Cerebral Cortex. 2001;11:298–311.

25. Wade A, Brewer A, Rieger J, Wandell B. Functional measurements of human ventral occipital cortex: Retinotopy and Color. Phil Trans R Soc Lond. 2002;357:963–973.

26. Gattass R, Sousa APB, Gross C. Visuotopic organization and extent of V3 and V4 of the Macaque. The Journal of Neuroscience. 1988;8:1831–45.

27. Goddard E, Mannion DJ, McDonald JS, Solomon SG, Clifford CWG. Color responsiveness argues against a dorsal component of human V4. Journal of Vision. 2011;11(4)(3):1–21. doi:10.1167/11.4.3.

28. Winawer J, Witthoft N. Human V4 and ventral occipital retinotopic maps. Visual Neuroscience. 2015;32:E020. doi:10.1017/S0952523815000176.

29. DeYoe EA, Bandettini P, Neitz J, Miller D, Winans P. Functional magnetic resonance imaging (FMRI) of the human brain. Journal of Neuroscience Methods. 1994;54:171–187.

30. Ogawa S, Tank DW, Menon R, Ellerman JM, Kim SG, Merkle H, et al. Intrinsic signal changes accompanying sensory stimulation: Functional brain mapping with magnetic resonance imaging. Proceedings of the National Academy of Sciences. 1992;89(13):5951–5955.

31. Puckett A, Mathis J, DeYoe E. An Investigation of Positive and Inverted Hemodynamic Response Functions Across Multiple Visual Areas. Human Brain Mapping. 2014;35(11):5550–5564.

32. Schira M, Wade A, Tyler C. Two-Dimensional Mapping of the Central and Parafoveal Visual Field to Human Visual Cortex. Journal of Neurophysiology. 2007;97:4284–4295.

33. Schira MM, Tyler CW, Breakspear M, Spehar B. The Foveal Confluence in Human Visual Cortex. Journal of Neuroscience. 2009;29(28):9050–9058. doi:10.1523/JNEUROSCI.1760-09.2009.

34. Dumoulin SO, Wandell BA. Population receptive field estimates in human visual cortex. NeuroImage. 2008;39(2):647–660. doi:10.1016/j.neuroimage.2007.09.034.

35. Brainard DH. The Psychophysics Toolbox. Spatial Vision. 1997;10:433–436.

36. Pelli D. The VideoToolbox software for visual psychophysics: Transforming numbers into movies. Multisensory research. 1997;10(4):437–442.

37. Fischl B, Salat DH, Busa E, Albert M, Dieterich M, Haselgrove C, et al. Whole Brain Segmentation: Automated Labeling of Neuroanatomical Structures in the Human Brain. Neuron. 2002;33(3):341–355. doi:10.1016/S0896-6273(02)00569-X.

38. Fischl B, Salat DH, van der Kouwe ajw, Makris N, Segonne F, Quinn BT, et al. Sequence-independent segmentation of magnetic resonance images. NeuroImage. 2004;23(Supplement 1):S69 – S84. doi:10.1016/j.neuroimage.2004.07.016.

39. Yoo T, Ackerman MJ, Lorensen WE, Schroeder W, Chalana V, Aylward S, et al. Engineering and Algorithm Design for an Image Processing API: A Technical Report on ITK - The Insight Toolkit. Studies in Health Technology and Informatics. 2002;85:586–592.

40. Friston KJ, Holmes AP, Worsley KJ, Poline JP, Frith CD, Frackowiak RSJ. Statistical parametric maps in functional imaging: A general linear approach. Human Brain Mapping. 1994;2(4):189–210. doi:10.1002/hbm.460020402.

41. Cox RW. AFNI: Software for analysis and visualization of functional magnetic resonance neuroimages. Comput Biomed Res. 1996;29:162–173.

42. Witthoft N, Nguyen ML, Golarai G, Larocque KF, Liberman A, Smith ME, et al. Where Is Human V4? Predicting the Location of hV4 and VO1 from Cortical Folding. Cerebral Cortex. 2013;9:2401–2408.

43. Smith A, Singh K, Greenlee M. Attentional Suppression of activity in the human visual cortex. Neuroreport. 2000;11(2):271–277.

44. Smith AT, Williams AL, Singh KD. Negative BOLD in the visual cortex: Evidence against blood stealing. Human Brain Mapping. 2004;21(4):213–220. doi:10.1002/hbm.20017.

45. Wade AR, Rowland J. Early Suppressive Mechanisms and the Negative Blood Oxygenation Level-Dependent Response in Human Visual Cortex. Journal of Neuroscience. 2010;30(14):5008–5019. doi:10.1523/JNEUROSCI.6260-09.2010.

46. IBM Corp. Released 2013. IBM SPSS Statistics for Mac, Version 22.0.Armonk, NY;IBM Corp.

47. Van Essen DC, Drury H, Dickson J, Harwell J, Hanlon D, Anderson CH. An Integrated Software Suite for Surface-based Analyses of Cerebral Cortex. Journal of the American Medical Informatics Association. 2001;8(5):443–459.

48. Fischl B, Dale AM. Measuring the thickness of the human cerebral cortex from magnetic resonance images. Proceedings of the National Academy of Sciences. 2000;97(20):11050–11055.

49. Saad ZS, Ropella KM, Cox RW, DeYoe EA. Analysis and use of FMRI response delays. Human Brain Mapping. 2001;13(2):74–93.

50. Polimeni J, Fischl B, Greve D, Lawrence W. Laminar analysis of 7 T BOLD using an imposed spatial activation pattern in human V1. NeuroImage. 2010;52:1334–1346.

51. Duvernoy HM, Delon S, Vannson JL. Cortical Blood Vessels of the Human Brain. Brain Research Bulletin. 1981;7:519–579.

52. Donahue MJ, Strother MK, Lindsey KP, Hocke LM, Tong Y, deB Frederick B. Time delay processing of hypercapnic fMRI allows quantitative parameterization of cerebrovascular reactivity and blood flow delays. Journal of Cerebral Blood Flow & Metabolism. 2016;36(10):1767–1779.

53. Krejza J, Mariak Z, Walecki J, Szydlik P, Lewko J, Ustymowicz A. Transcranial color Doppler sonography of basal cerebral arteries in 182 healthy subjects: Age and sex variability and normal reference values for blood flow parameters. American Journal of Roentgenology. 1999;172:213–8.

54. Bisaria KK. Anatomic variations of venous sinuses in the region of the torcular Herophili. Journal of Neurosurgery. 1985;62(1):90–95. doi:10.3171/jns.1985.62.1.0090.

55. Saiki K, Tsurumoto T, Okamoto K, Wakebe T. Relation between bilateral differences in internal jugular vein caliber and flow patterns of dural venous sinuses. Anatomical Science International. 2013;88(3):141–150. doi:10.1007/s12565-013-0176-z.

56. Zeki S. Improbable areas in the visual brain. Trends in Neurosciences. 2003;26:23–6.

57. Jezzard P, Balaban R. Correction for Geometric Distortion in Echo Planar Images from B0 Field Variations. Magnetic Resonance in Medicine. 1995;34(1):65–73.

58. Jezzard P, Clare S. Sources of Distortion in Functional MRI Data. Human Brain Mapping. 1999;8:80–85.

59. Pasley BN, Inglis BA, Freeman RD. Analysis of oxygen metabolism implies a neural origin for the negative BOLD response in human visual cortex. NeuroImage. 2007;36(2):269–276. doi:10.1016/j.neuroimage.2006.09.015.

